# Arabidopsis latent virus 1, a comovirus widely spread in *Arabidopsis thaliana* collections

**DOI:** 10.1101/2022.07.21.500942

**Authors:** Ava Verhoeven, Karen J. Kloth, Anne Kupczok, Geert H. Oymans, Janna Damen, Karin Rijnsburger, Zhang Jiang, Cas Deelen, Rashmi Sasidharan, Martijn van Zanten, René A.A. van der Vlugt

## Abstract

Transcriptome studies of Illumina RNA-seq datasets of different *Arabidopsis thaliana* natural accessions and T-DNA mutants revealed the presence of two virus-like RNA sequences which showed the typical two segmented genome characteristics of a comovirus.
This comovirus did not induce any visible symptoms in infected Arabidopsis plants cultivated under standard laboratory conditions. Hence it was named Arabidopsis latent virus 1 (ArLV1). Virus infectivity in Arabidopsis plants was confirmed by RT-qPCR, transmission electron microscopy and mechanical inoculation. ArLV1 can also mechanically infect *Nicotiana benthamiana,* causing distinct mosaic symptoms.
A bioinformatics investigation of Arabidopsis RNA-Seq repositories, including nearly 6500 Sequence Read Archives (SRAs) in the NCBI SRA database, revealed the presence of ArLV1 in 25% of all archived natural Arabidopsis accessions and in 8.5% of all analyzed SRAs. ArLV1 could also be detected in Arabidopsis plants collected from the wild.
ArLV1 is highly seed-transmissible with up to 40% incidence on the progeny derived from infected Arabidopsis plants. This has likely led to a worldwide distribution in the model plant Arabidopsis with yet unknown effects on plant performance in a substantial number of studies.

**Plain language summary:** We identified Arabidopsis latent virus 1 (ArLV1), a comovirus that infects the model plant *Arabidopsis thaliana* without causing any visible symptoms. It is efficiently spread by transmission via seeds to the plant progeny. ArLV1 is infectious to Arabidopsis plants and another model plant, *Nicotiana benthamiana.* By analyzing public sequencing data, we found that ArLV1 is widely spread in Arabidopsis laboratory collections worldwide. Moreover, it was also detected in wild Arabidopsis plants collected from different locations in the Netherlands and Spain, suggesting that it is a virus that naturally occurs in Arabidopsis.

## Introduction

Many plant biology studies involve *Arabidopsis thaliana* as model plant. Although community-serving Arabidopsis stock centers, such as the Arabidopsis Biological Resource Centre (ABRC) (Scholl & Anderson, 1994), test for seed-borne diseases, usually only visual detection methods and germination tests are applied (Rivero *et al*., 2014). Moreover, few research labs have reported to test their seed stocks for known virus contaminations before running experiments. The *Arabidopsis thaliana* genome sequence became publicly available in 2000 (The Arabidopsis Genome Initiative, 2000). Since then, whole transcriptome sequencing has become one of the most common tools to decipher plant physiological processes. This untargeted approach can also reveal unexpected presence of other biological agents and provides information about possible unnoted infections and contaminations (Villamor *et al*., 2019).

Particularly viruses can be hiding in plants and in seed material (Cobos *et al*., 2019). The majority of well-studied viruses cause disease symptoms in agriculturally important crops with sometimes severe effects on plant morphology, physiology and yield (Prasad *et al*., 2020). In nature however, plants are often infected with viruses that do not cause any apparent disease symptoms, so-called latent infections (Shates *et al*., 2019). Many viruses may even be beneficial to their hosts in a mutualistic symbiosis (Roossinck, 2011). Extending the knowledge on these latent viruses will contribute to the beneficial exploitation of viruses in cultivated crops (Takahashi *et al*., 2019).

Here, we describe a comovirus; Arabidopsis latent virus 1 (ArLV1), that we encountered in Arabidopsis RNA sequencing datasets generated in our labs and which was found to be widespread in other datasets obtained from sequence data repositories. We found that plants from several Arabidopsis accessions, including the widely used accession Col-0 (CS60000), tested positive for ArLV1. We identified different isolates of the virus across the NCBI Sequence Read Archives and investigated disease symptoms, infectivity, plant growth and effects on the Arabidopsis transcriptome and on abiotic stress resilience. Regular screening for the presence of this widely present – but unnoted – virus and further investigation of its possible effects is highly relevant for the plant science community working with Arabidopsis.

## Materials and methods

### ArLV1 identification and genome assembly

The RNA-Seq dataset of Kloth *et al.* (2016) was mapped against the TAIR10 Arabidopsis reference transcriptome (Lamesch *et al*., 2011) with Tophat version 2.0.13 and intron length 20-2000. The dataset from Utrecht University was mapped against the Araport10 reference transcriptome with *KALLISTO* (Bray *et al*., 2016) (Supporting Information Methods S1). Un-mapped reads were *de novo* assembled in CLC Genomic Workbench V9 (Qiagen, Aarhus A/S Denmark) using standard settings and resulting contigs were blasted against the NCBI RefSeq database. Contigs showing a clear identity to different comoviruses were retained for further analysis. Per Arabidopsis genotype (ALL1-3, Pent-1, Ep-0 and Col-0), we assembled unique contigs, of which the virus isolate in ALL1-3 from the Wageningen dataset was submitted to GenBank under accession numbers MH899120.1 (RNA1) and MH899121.1 (RNA2) respectively. This Wageningen isolate is later referred to as ArLV1_A, the Col-0 isolate from Utrecht as ArLV1_B. To identify the phylogenetic position of this virus within in the genus *Comovirus,* RefSeq comovirus sequences were downloaded from NCBI and the amino acid sequence of the conserved Pro-Pol region from RNA1 was aligned using MAFFT v7.475 with the auto option (Katoh & Standley, 2013). Maximum likelihood (ML) phylogeny was reconstructed using IQ-TREE v2.0.3 and 1000 ultrafast bootstrap replicates (Minh *et al*., 2020).

### Datamining of NCBI Sequence Read Archives

Illumina-generated *Arabidopsis thaliana* RNA-Seq datasets (SRAs) were downloaded from the NCBI Sequence Read Archive (https://www.ncbi.nlm.nih.gov/sra) and searched for the presence of ArLV1 RNA1 and RNA2 sequences with the blastn_vdb program from the NCBI SRA toolkit version 2.9.0 (min. 500 reads per SRA dataset; automatization script available on Zenodo).

### Phylogenetic analysis

From the NCBI SRA output, accessions containing ArLV1 were selected based on the presence of at least 500 reads of RNA2, representing full coverage. From these, one SRA each from 38 randomly chosen accessions, was selected from which the consensus nucleotide sequences of RNA1 and RNA2 were retrieved by Reference Mapping (CLC Genomic Workbench v20) against the ArLV1 sequences (MH899120.1 (RNA1) and MH899121.1 (RNA2), respectively) with options ‘Low coverage definition threshold = 3’ and ‘insert N-ambiguity symbol’. ML phylogenies were reconstructed using the conserved Pro-Pol region from RNA1 (Le Gall et al., 2008) and ORF2 from RNA2 of these 38 consensus sequences together with four sequences from different Arabidopsis ecotypes from Wageningen and Utrecht (ALL1-3_Wageningen, Pent-1_Wageningen, Ep-0_Wageningen and Col-0_Utrecht) using IQ-TREE v2.0.3 and 1000 ultrafast bootstrap replicates (Minh et al., 2020). Evidence of recombination was assessed using the Phi test implemented in SplitsTree (Huson & Bryant, 2006; Bruen et al., 2006). A worldmap of ArLV1 occurrences, based on available GPS data (https://1001genomes.org) for 36 accessions with available latitude and longitude coordinates, was made with the R package ggplot2.

### ArLV1 inoculations

ArLV1 inoculum was obtained by harvesting leaf material from ArLV1 positive Arabidopsis plants and grinding it (1:1 w/v) in an inoculation buffer (0.03 M phosphate buffer, pH of 7.2). For inoculation, leaves of Arabidopsis plants in leaf stage 8-10, or *Nicotiana benthamiana* plants in leaf stage 5-8 were lightly dusted with carborundum powder and inoculum was applied by gentle rubbing. Plants were rinsed with water 10 minutes after inoculation. Virus symptoms were assessed visually 7-10 days post inoculation (DPI). Presence of virus in inoculated plants was checked by SYBR Green RT-PCR and transmission electron-microscopy (TEM) with a JEOL, JEM1400 Plus using a leaf dip assay according to standard protocols (Hayat & Miller, 1990).

### Virus detection by reverse transcriptase (RT), real-time quantitative PCR (RT-qPCR)

For a detailed step-by-step protocol of how samples were collected and used in RT-qPCR, please refer to Supporting Information Fig. S1. In addition to this protocol, at least two primer pairs were used per sample; one for RNA1 (RNA1_1; ArLV-RNA1-Fw: TCTGCCAGTACTGGAGAGG and ArLV1-RNA1-Rv: GTCATCCAACAAATAGGAAC) and one for RNA2 (RNA2; ArLV1-RNA2-Fw: CACCAATAACACCCCAAAA and ArLV1-RNA2-Rv: GCATTTCCACAGAGTCTCG). For the seed transmission experiments, an additional primer pair for RNA1 was used (RNA1_2; ArLV1-RNA1-Fw: TGTCGTGATAACTGATGG and ArLV1-RNA1-Rv: CTAACCTCTTTCCTCCCC). dCT values were calculated per primer pair by subtracting the average control CT values from the same RT-qPCR run. If multiple primer pairs for RNA1 were used, the average delta CT of the two was calculated and used in visualization. *K-means* clustering was used to separate the positive samples from the negative samples.

## Results

### Identification of Arabidopsis latent virus 1 (ArLV1)

An analysis of RNAseq transcriptome datasets from several natural Arabidopsis accessions, of which some were part of a study by Kloth et al. (2016), hereafter referred to as the ‘Wageningen’ dataset, showed for some samples an unexpectedly low mapping percentage of plant reads (as low as 17.4%) in the alignments to the TAIR10 Arabidopsis reference transcriptome (Lamesch *et al*., 2011) (Fig. **1a**). *De novo* assembly of the unmapped reads identified two contigs with respective lengths of 5953 and 3600 nucleotides, displaying a segmented genome organization typical for viruses from the genus *Comovirus,* family *Secoviridae,* order *Picornavirales* (Thompson *et al*., 2017) (Fig. **1b**). To assess the phylogenetic position of this comovirus, we compared the amino acid sequence of the highly conserved Pro-Pol region of RNA1 with the Pro-Pol region of other comoviruses (Le Gall *et al*., 2008). The highest level of nucleotide identity of this comovirus is 68% with *Radish mosaic virus* (RaMV; NC_010709). This analysis clearly identified the virus as a distinct comovirus (Fig. **1c**). Samples of two Arabidopsis accessions contained extremely high numbers of reads from the newly identified comovirus: accession ALL1-3 (CS76090) up to 88.2% (isolate ArLV1_A), and accession Pent-1 (CS76209) up to 83.0%. We also identified an extremely high abundance of similar RNA1 and RNA2 reads from the same comovirus in some, but not all, RNA-sequenced samples of accession Col-0 at Utrecht University with a mapping percentage of up to 90.08% of viral reads (isolate ArLV1_B) (Fig. **1d**). The occurrence of the virus in samples was not linked to the applied treatments (neither abiotic stress or aphids). As this comovirus does not seem to cause any apparent visible symptoms in any of the Arabidopsis accessions in our studies, we named the virus Arabidopsis latent virus 1 (ArLV1).

**Figure 1.**
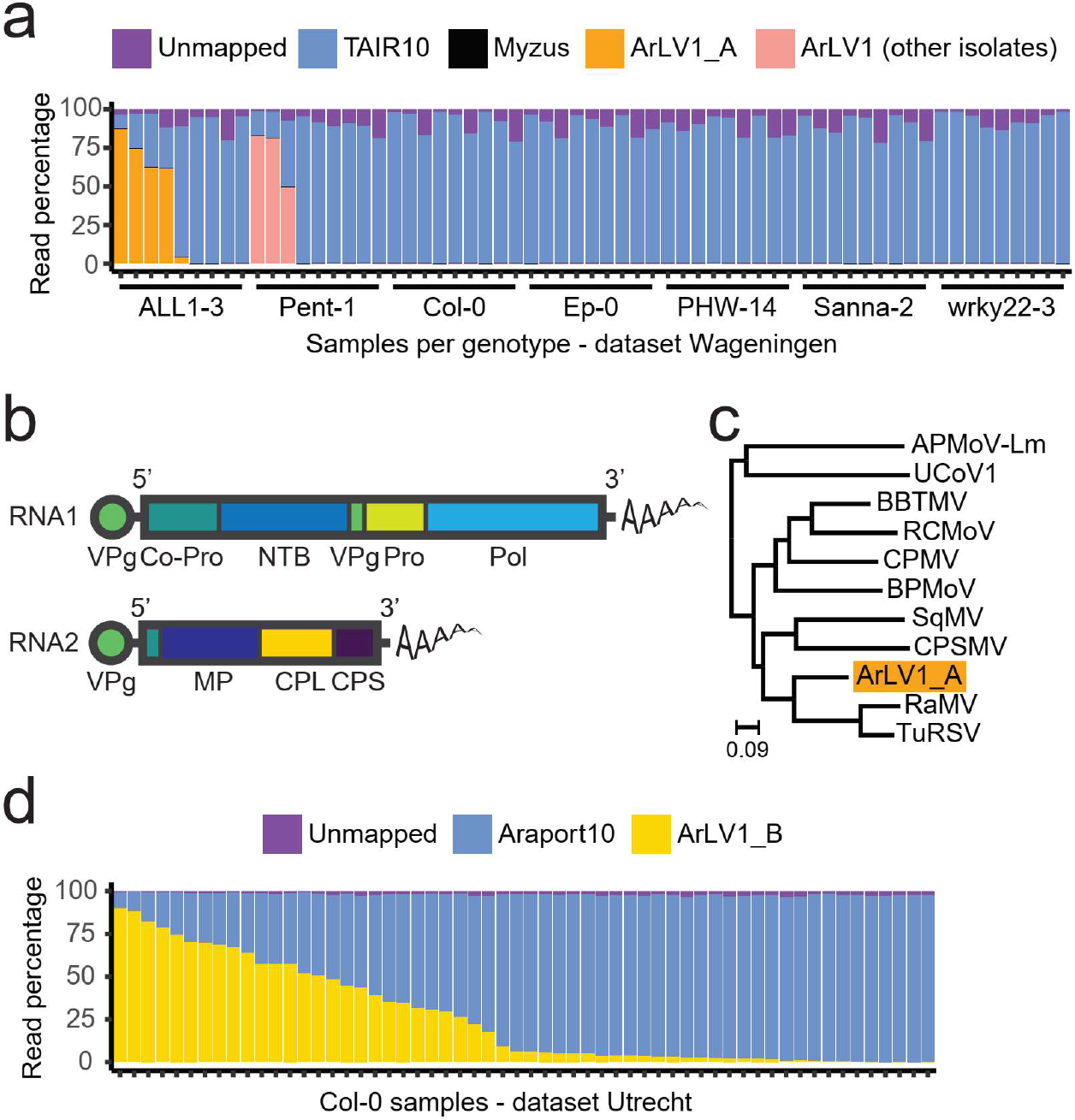
Occurrence and identification of ArLV1 in Arabidopsis. (**a**) Mapping percentages of the reads in the Wageningen RNA-Seq dataset. As this dataset involved leaf material from both naïve and aphid-infested samples (Kloth *et al,.* 2016), reads of *Myzus persicae* aphids were included in our analysis as well, but only few were identified. Sequence information for further analyses was obtained from ALL1-3, and the virus isolate was named ArLV1_A. (**b**) Schematic representation of RNA1 and RNA2 of a typical comovirus, adapted from (King *et al*., 2012). VPg: genome-linked viral protein; Co-Pro: proteinase cofactor; NTB: NTP-binding proteins; Pro: proteinase; Pol: RNA-dependent RNA polymerase; MP: movement protein; CPL and CPS: large and small capsid proteins. (**c**) Maximum likelihood (ML) tree of the translated Pro-Pol region from RNA1 of ArLV1_A (the isolate from ALL1-3) and ten other comoviruses: APMoV-Lm (andean potato mottle virus; MN176101), UCoV1 (Ullucus virus C; MH645163), BBTMV (broad bean true mosaic virus; NC_022004), RCMoV (red clover mottle virus; NC_003741), CPMV (cowpea mosaic virus; NC_003549), BPMoV (bean pod mottle virus; NC_003496), SqMV (squash mosaic virus; NC_003799), CPSMV (cowpea severe mosaic virus; NC_003545), RaMV (radish mosaic virus; NC_010709), TuRSV (turnip ringspot virus; NC_013218). Branch lengths (scale) represent amino acid substitutions per site. Note that this is an unrooted tree. (**d**) Mapping percentages of the ArLV1_B reads in the Utrecht RNA-Seq dataset from Arabidopsis accession Col-0, involving leaf samples from an abiotic stress experiment (combinations of mild drought, high temperature and submergence; see Supporting Information Methods S1 and Morales *et al*., (2022)).

### ArLV1 detection, inoculation and transmission

To assess possible plant infections, we developed and validated a RT-qPCR suitable for plant leaf material (Supplemental Information Fig. S1). ArLV1 isolates could be mechanically inoculated from infected Arabidopsis plants, grown from infected seed batches, to *N. benthamiana,* known for its susceptibility to many plant viruses (Goodin *et al*., 2008) and healthy Arabidopsis plants. *N. benthamiana* plants showed symptoms of leaf mottling and mosaic patterns at 5-7 DPI (Fig. **2a**) and all inoculated plants tested positive when compared with mock-inoculated plants (Fig. **2b**). Arabidopsis plants infected with ArLV1 never showed any visible symptoms and could not be visually distinguished from healthy plants. However, transmission electron microscopy of ArLV1 infected Arabidopsis leaf extracts did show typical comovirus particles (Fig. **2c**). We used *N. benthamiana* leaf tissue to mechanically inoculate Arabidopsis Col-0 with the isolates ArLV1_A and ArLV1_B with an efficiency of 88% (79/90) (Supporting Information Fig. S2). Taken together, we can state that ArLV1 can be mechanically transferred and has the ability to infect both *N. benthamiana* and Arabidopsis. As some comoviruses can be efficiently transmitted via seeds (Gergerich & Scott, 1996), we tested the progeny of four different ArLV1-infected Arabidopsis Col-0 parent plants for presence of ArLV1. In total, 39.1% of the 46 plants grown from these seed batches tested positive for ArLV1 (Fig. **2d**, Supporting Information Fig. S3), indicating seed transmission of ArLV1.

**Figure 2.**
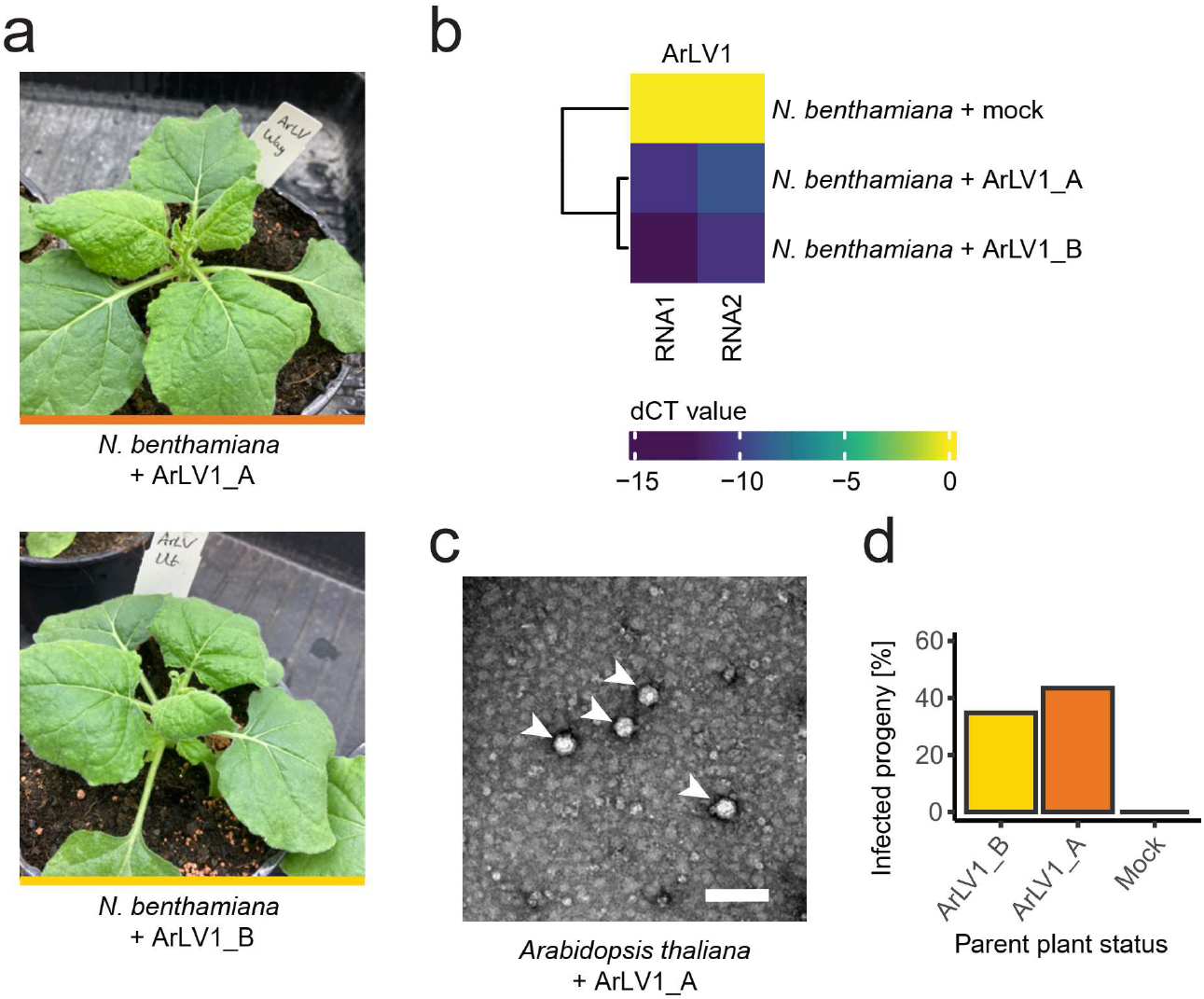
Mechanical inoculation, detection and seed transmission of ArLV1. (**a**) *N. benthamiana* plants at two weeks after mechanical inoculation with two isolates or ArLV1: ArLV1_A (upper photo) and ArLV1_B (lower photo), both showing ArLV1-induced leaf mottling symptoms. (**b**) RT-qPCR detection of RNA1 and RNA2. This experiment was repeated multiple times with comparable results. (**c**) ArLV1_A in infected Arabidopsis leaves, visualized by transmission electron microscopy. Arrowheads indicate the viral particles (scale bar is 100 nm). (**d**) The percentage of infected progeny from Arabidopsis Col-0 parent plants infected with the isolate ArLV1_A or the ArLV1_B. Infection was detected by RT-qPCR on leaf material (Supporting Information Fig. S1 and S2).

### Occurrence and phylogeny of ArLV1

To reveal if ArLV1 is also present in Arabidopsis datasets other than the Wageningen and Utrecht datasets, we initiated a search in publicly available Arabidopsis RNA-Seq datasets. Out of a total of 6477 RNA-Seq datasets analyzed, 547 (8.45%) contained at least 500 reads mapping to RNA1 and 500 reads mapping to RNA2 of ArLV1, indicating a full coverage of the viral genome. Altogether, ArLV1 was detected in 176 out of 711 accessions (24.75%) in the public datasets analyzed. To assess the genetic diversity of ArLV1, nucleotide sequences from the Pro-Pol regions of RNA1 and from the full ORF of RNA2 of four isolates from different Arabidopsis ecotypes from Wageningen and Utrecht and 38 different NCBI-derived datasets, each with a full coverage of RNA1 and RNA2, were used for phylogenetic analyses (Fig. **3a**; Supporting Information Fig. S5, S6). The resolved phylogenetic trees supported a separation of isolates in three distinct clades. Clade 1 was represented by 55% of the isolates and occurred across different continents, while clade 2 and 3 were represented by 29% and 17% of the isolates, respectively, and originated from Eurasia, except from 1 isolate of clade 2 from the US (Pent-23) (Fig. **3b**; Supporting Information Table S1). The isolates within clade 1 and 2 were highly similar while six isolates in the third clade were more divergent (Supporting Information Fig. S5, S6). Notably, the phylogenies showed signs of reassortment (e.g., accessions Fell2-4 and Pent-23 fall in clade 1 for RNA1 and in clade 3 and clade 2, respectively, for RNA2) (Supporting Information Fig. S5, S6). We also of detected recombination within each alignment (Phi test p-value<0.01 for each alignment). To find out if ArLV1 was also present in wild Arabidopsis populations, we sampled 27 different plants in Arnhem, Wageningen and Woerden (The Netherlands), where presence of ArLV1 outside of the laboratory environment was confirmed in independent RT-PCR assays for 8/27 plants (Supporting Information Fig. S4). In addition, ArLV1-presence in wild Arabidopsis plants from Ciruelos de Coca and Carbonero (Spain) (Pagán *et al*., 2010) was confirmed by RT-PCR and Sanger sequencing confirmation (C. Carrasco-López & F. García Arenal, pers. comm.).

**Figure 3.**
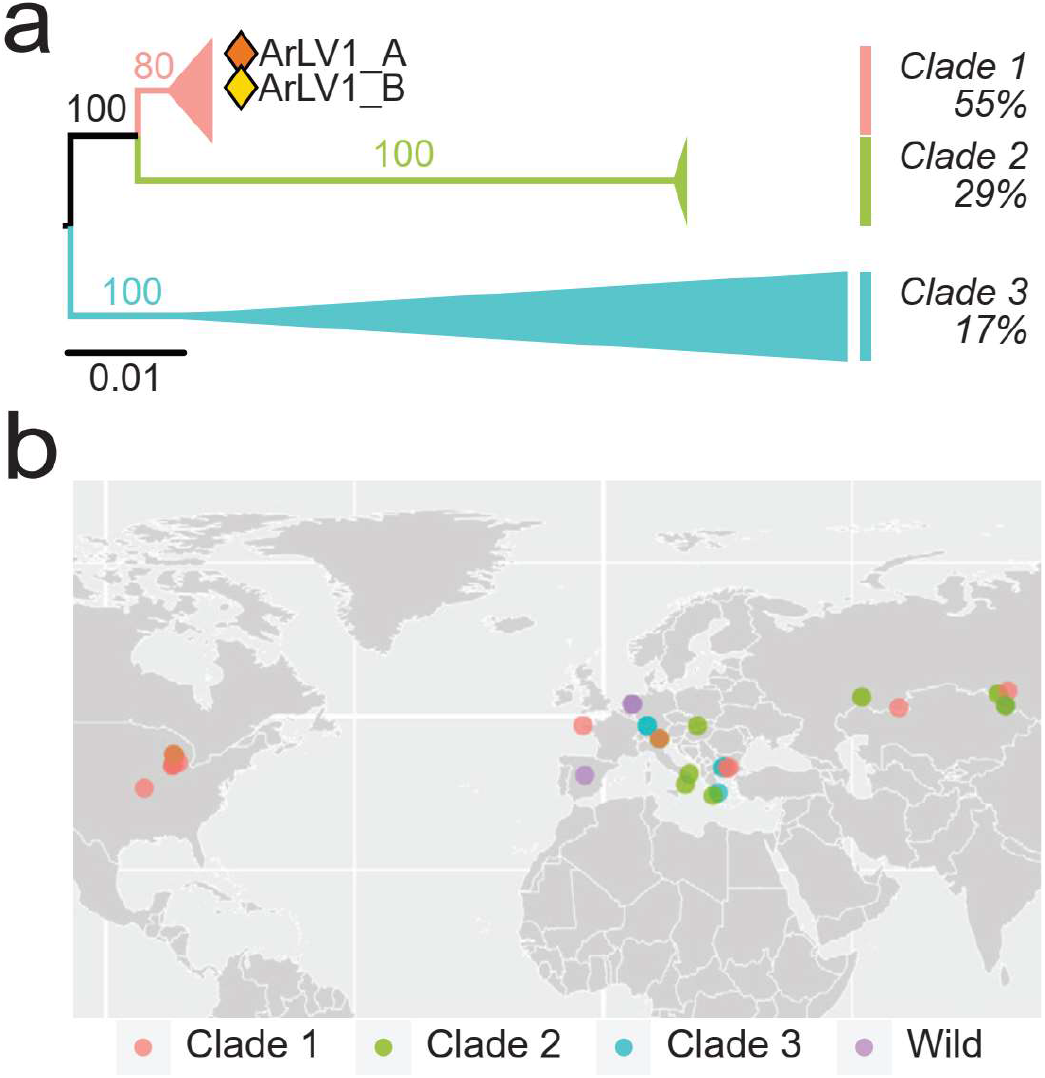
Phylogeny and geographical information of different ArLV1 isolates. (**a**) Collapsed maximum likelihood tree of the nucleotide sequences of ORF2 on RNA2 of 38 ArLV1 isolates. Branch lengths (scale) represent nucleotide substitutions per site. (**b**) World map with the coordinates of 36 different ArLV1 isolates, colored by their RNA2 clade. For the ArLV1 isolates collected outside of the laboratory environment (wild), we do not have clade data available. Genotypes from Arabidopsis accessions collected at relatively close proximity are represented by overlapping dots.

### Analysis of ArLV1 effects

Although ArLV1 produces no visible symptoms in Arabidopsis, we wanted to study possible effects that ArLV1-infection may have on the transcriptome responses of Arabidopsis plants. To this end, we compared the transcriptomes of seven samples from the Utrecht transcriptome analysis. This dataset included four samples with high numbers of viral reads (a mapping percentage of 78.94-90.08%) and three samples of plants of identical age and growth conditions, but with low numbers of viral reads (a mapping percentage of 0.01-9.56%). We did not find any significant differentially expressed Arabidopsis genes between the two groups (Fig. **4a**). However, we did observe a slight but significantly lower chlorophyll content in plants inoculated with ArLV1 compared to mock-inoculated plants, regardless of the availability of water (Fig. **4b**) (Supporting Information Methods S3). Other morphometric traits, such as leaf number or the leaf surface area were unaffected by the virus (Supporting Information Fig. S7). When watering was stopped to induce drought conditions, mock-inoculated plants wilted significantly faster than ArlV1-inoculated plants (Fig. **4c**). The same phenotype was observed for *N. benthamiana* plants subjected to similar drought conditions (Fig. **4d**). Next, we quantified the area of the pectin mucilage layer present on the external surface of seeds. The presence of this layer is associated with seed longevity and the size and integrity is known to be altered by several viruses (Bueso *et al*., 2017) (Supporting Information Methods S3). Although the different growing conditions in Wageningen and Utrecht significantly (P=0.004) affected the area of the pectin mucilage layer, ArLV1-infection of the parent plant did not affect the size of the mucilage layer (P=0.147) (Supporting Information Fig. S8a). Likewise, Arabidopsis Col-0 plants inoculated with the two isolates of ArLV1 or mock-inoculated did not differ from each other in the onset of flowering (Supporting Information Fig. S8b), suggesting that ArLV1 is not affecting these fitness parameters. Taken together, these results indicate that while ArLV1 is a largely latent virus to Arabidopsis, some minor phenotypic effects can be observed such as lower chlorophyll content and improved drought tolerance. The latter effect was observed in both Arabidopsis and *N. benthamiana*.

**Figure 4.**
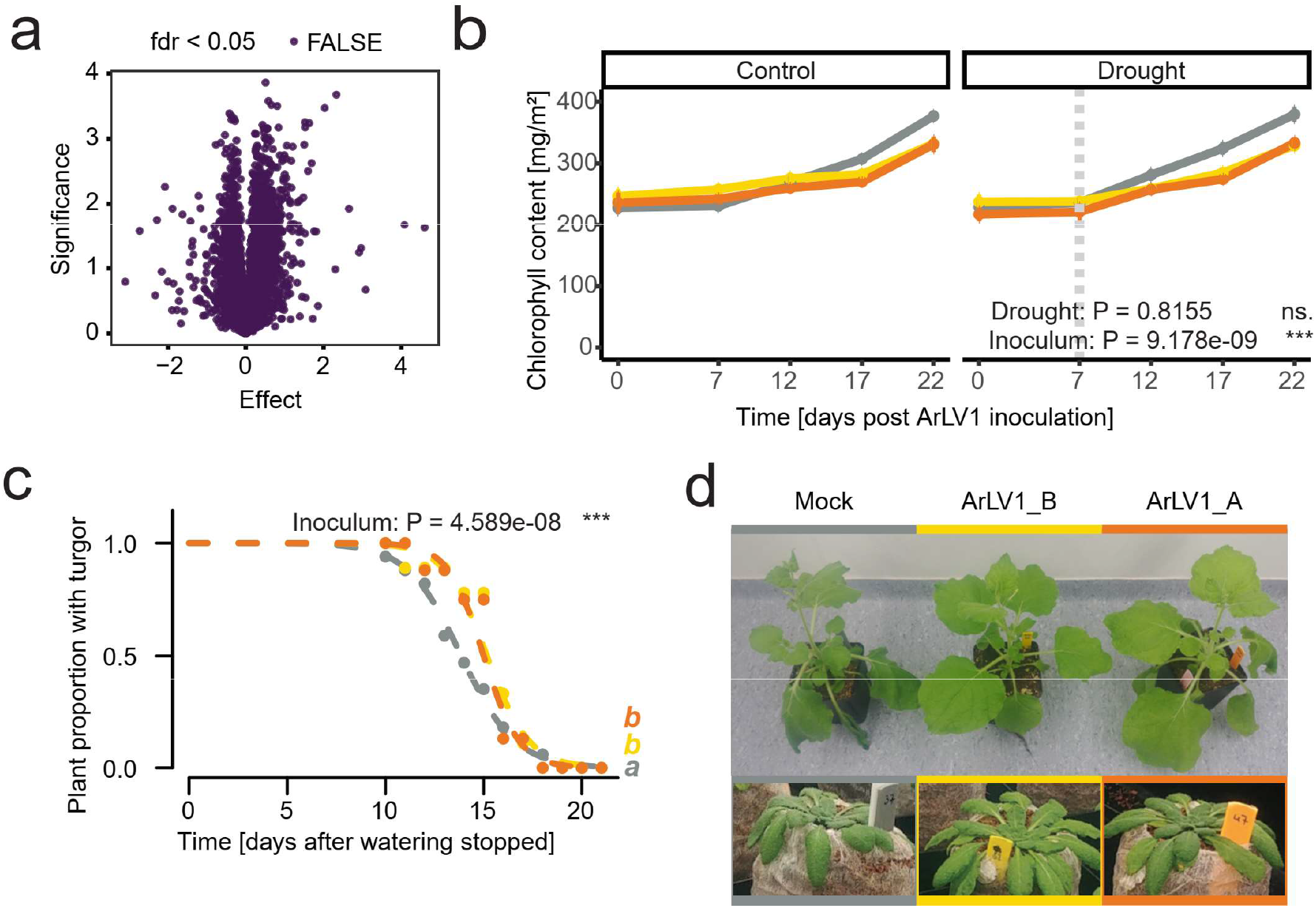
Transcriptomic and phenotypic effects of ArLV1. (**a**) Arabidopsis Col-0 transcriptome analysis of four samples with high ArLV1_B (Utrecht isolate) read mapping (average = 81.6%) compared with three samples with low ArLV1_B read mapping (average = 4.18%). (**b**) Chlorophyll content levels of the 8^th^ true leaf, following ArLV1 infection in plants subjected to drought (right panel) or kept in well-watered control conditions (left panel). Inoculation with ArLV1_A (orange), ArLV1_B (yellow) or Mock (grey) occurred at Time = 0 DPI and drought was applied at Time = 7 DPI; vertical dotted bar; see Supporting Information Methods S1. (**c**) Survival curve of the fraction of plants that maintained turgor after watering was stopped (at Time = 0 days). The experiments (**b, c**) were repeated twice with similar results (n>20. Both repeats were included in the statistical analysis using (generalized) mixed linear models with repeats as a random variable and an alpha of 0.05 (different letters (a, b) represent significant differences). The p-values of Inoculum or Drought represents the effect of inoculation with either ArLV1_A, ArLV1_B or Mock, or the effect of the application of drought. (**d**) Image showing representative *N. benthamiana* and Arabidopsis plants inoculated with ArLV1_A, ArLV1_B or Mock displaying wilting after four days (N. *benthamiana)* or 15 days (Arabidopsis) of water deprivation.

## Discussion

The advent of high throughput sequencing technologies has greatly contributed to the understanding of the concept of holobionts (Nobori *et al.*, 2018), which includes not only the study organism itself but also its associated communities (Hassani *et al*., 2018). In contrast to the use of microarrays, high throughput sequencing can reveal the presence of unexpected and unrevealed organisms and biological agents in study systems (Vandenkoornhuyse *et al*., 2015), especially for viruses (Massart *et al*., 2014; Maclot *et al*., 2020). As sequencing has become more affordable and prevalent, opportunities arise to uncover the unknown metagenomes of our study systems. This may reveal critical microbial factors that potentially influence plant (physiological) processes, and, therefore, urges us to investigate unexpected results, such as illustrated here.

In this study, Illumina-derived RNA-seq data sets from Arabidopsis transcriptome studies with exceptionally low numbers of plant-specific reads revealed the presence of a so far uncharacterized plant virus belonging to the genus *Comovirus.* Given its apparent latent nature in Arabidopsis we named this virus Arabidopsis latent virus 1 (ArLV1). We showed the infectivity of this newly discovered virus for Arabidopsis and *N. benthamiana* and its high transmissibility to Arabidopsis progeny via seeds from ArLV1 infected plants. TEM studies confirmed typical comovirus particles in infected Arabidopsis plants and a RT-qPCR test was developed that allows detection of the virus in plant material.

From the nearly 6500 public Arabidopsis sequence read archives that were tested, 8.45% contained evidence of an ArLV1 infection, accounting for near 25% of the ‘natural’ Arabidopsis accessions and some mutant lines in the dataset. This indicates that ArLV1 is present in the Arabidopsis stocks of laboratories worldwide. We also confirmed its presence in wild Arabidopsis populations, both in The Netherlands and in Spain. 38 Arabidopsis accessions were selected for a phylogenetic analysis of ArLV1 sequences, including the widely used accession Col-0. The phylogenetic analysis divided these virus sequences into three clades with different geographical distributions, where occasional recombination and re-assortment also occurs. This, as well as its apparent latent nature, its occurrence in plants directly collected from the wild and a large number of laboratory stocks of Arabidopsis accessions collected from many geographical regions, clearly suggest that ArLV1 is a virus that has been naturally associated with Arabidopsis for a long period of time.

Our RNA-Seq datasets obtained from different Arabidopsis accessions in independent studies both in Wageningen and Utrecht show that an ArLV1 infection in Arabidopsis can result in more than 90% of virus-specific reads. This is an indication that ArLV1 can potentially reach very high titers in infected plants. Interestingly, large differences in ArLV1-specific read numbers (both absolute and relative to the total read count) were observed in datasets from Wageningen and Utrecht, between plants from the same accession grown in the same experiment. The two labs in Wageningen and Utrecht used different plant growth conditions (Supporting Information Methods S1) making it unlikely that specific growing conditions are related to these differences. The reason(s) for the high variation in ArLV1 virus titers remain to be elucidated. Although we did not identify public data sets with high amounts of virus-specific reads, we suspect that unpublished Arabidopsis transcriptome studies possibly also contain comparable high numbers of ArLV1 reads, but have not been reported and made available due to the low number of plant-specific reads.

Although a direct comparison of the transcriptomes of Arabidopsis plants with high and low viral infections did not reveal differentially expressed genes, ArLV1 did affect drought resilience, putatively via virus-induced reduced stomatal conductance (Pasin *et al*., 2020; Manacorda *et al*., 2021). Next to the obvious negative impacts on plant morphology, physiology and yield (Prasad *et al*., 2020), viruses are also known to affect their host positively in their tolerance to abiotic stress (Gorovits *et al*., 2019; Rahman *et al*., 2021; Aguilar & Lozano-Duran, 2022). Some viruses even change from parasitic to mutualistic with a change of environmental conditions (González *et al*., 2021). The protective effect of plant viruses is mainly observed for drought (Mishra *et al*., 2021), but some studies also report that viruses can have a positive effect on plant responses to (other) abiotic stresses, such as high temperature (Anfoka *et al*., 2016) or salt stress (Sinha *et al*., 2021). To our knowledge this is a first example of a comovirus, exerting such a positive effect. Further research is needed to reveal the underlying molecular mechanisms of how ArLV1 can affect plants tolerance to drought, and if similar effects can be observed for other abiotic stressors.

Taken together, we have identified a thus far unknown comovirus that is widely distributed in *Arabidopsis thaliana* in laboratories and the wild. The high prevalence in transcriptome datasets and its high potential for seed transmission make it safe to assume that ArLV1 will be present in research set-ups with the model species Arabidopsis with a significant part of the plants within a given experiment unknowingly infected with ArLV1. This may have unknown consequences for the interpretation of data obtained from these studies. Given its largely latent nature, ArLV1 has likely remained unnoticed for long and through its efficient seed transmission has spread worldwide. Its prevalence and lack of obvious disease symptoms, makes ArLV1 a plant virus that needs to be treated with scrutiny. We recommend routine screening to detect the presence of ArLV1 in seed stocks in labs and public repositories. The virus can be easily detected in plant samples via the qRT-PCR based method described in Supplementary Information Fig. S1. This will permit rapid selection of ArLV1-free plants and seed batches before proceeding with experiments, preventing potential confounding effects of the virus.

## Acknowledgments

This research was supported by the Netherlands Organization for Scientific Research (NWO), project numbers: 867.15.031, to A.V., R.S., M.v.Z., 016.VIDI.171.006 to R.S., a China Scholarship Council (CSC) project number 201806170025 to J.Z., The Graduate School Experimental Plant Sciences (project V-GENE) to A.K. and the NWO Technology Foundation Perspective Programme ‘Learning from Nature’ (STW10989) and the ZonMw programme ‘Enabling Technologies’ (435000017) to K.J.K. We wish to thank Emilyn Matsumura for critical reading of the manuscript, Rebecca Augsburger for technical assistance and Jan C. van Haarst for his help with the analysis of the initial dataset containing the virus. We thank Cristian Carrasco-López and Fernando García Arenal (CISC, Madrid) for the helpful discussions and sharing their unpublished data on the ArLV1-confirmation experiments performed on wild plants collected in Spain. The authors declare no competing interests.

## Author Contributions

A.V., K.J.K., and R.A.A.v.d.V. designed the experiments; A.V., K.J.K., R.A.A.v.d.V., A.K., G.O., K.R., J.Z., J.D., and C.D. performed the experiments; A.V., K.J.K., and R.A.A.v.d.V. analyzed the results. A.V., K.J.K., R.A.A.v.d.V., R.S. and M.v.Z. wrote the paper.

## Data Availability

The sequences of the ArLV1_A isolate in ALL1-3 from the Wageningen dataset are available at NCBI (www.ncbi.nlm.nih.gov/) under accession numbers MH899120.1 (RNA1) and MH899121.1 (RNA2) respectively. The Python scripts used for ArLV1 detection in the NCBI-SRA have been deposited in Zenodo (https://zenodo.org/record/6834314#.YtA3ei8Rqys). RNA-Seq datasets were deposited at NCBI under project number PRJNA858638 (Wageningen dataset) and project number (pending) (Utrecht dataset). All other data are available from the corresponding author upon request.

