## Supporting Information for "Arabidopsis latent virus 1, a comovirus widely spread in *Arabidopsis thaliana* collections"

The following Supporting Information is available for this article:

**Figure S1** Protocol for ArLV1-detection in plant material

**Figure S2** Arabidopsis inoculation efficiency

**Figure S3** Arabidopsis seed transmission of ArLV1

**Figure S4** ArLV1 RNA2 PCR fragments of wild collected Arabidopsis

**Figure S5** Phylogenetic tree of the Pro-Pol region from ArLV1 RNA1

**Figure S6** Phylogenetic tree of the full ORF from ArLV1 RNA2

**Figure S7** Leaf number and leaf surface area of plants subjected to ArLV1-inoculation

**Figure S8** Seed mucilage layer and flowering time are not affected by ArLV1

**Table S1** Arabidopsis SRA-datasets included in the geographical analysis

**Methods S1** RNA-Seq approaches and plant growth, treatment and sampling conditions

**Methods S2** Transcriptomic analysis of Arabidopsis Col-0 accession

**Methods S3** Phenotypic parameters

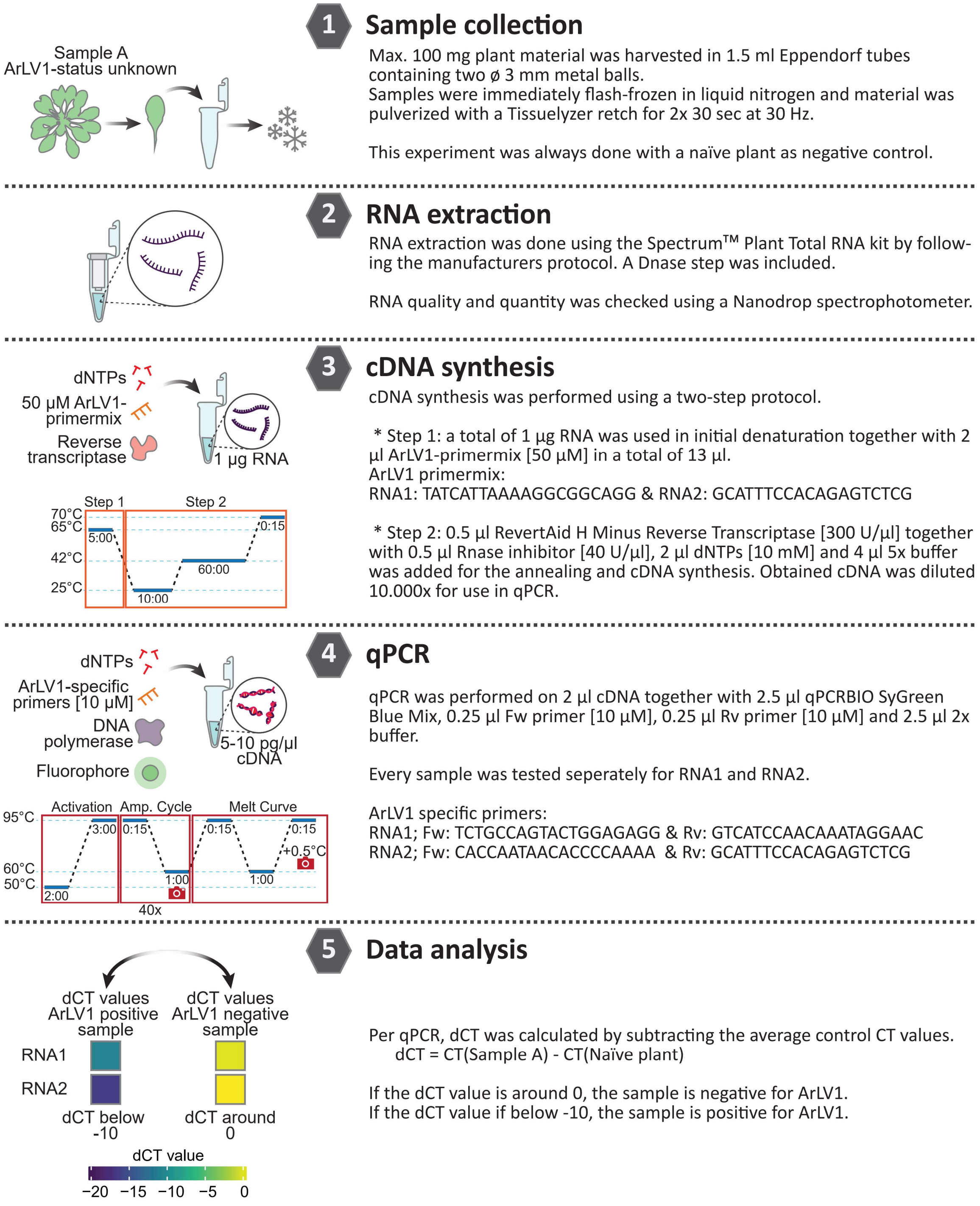


**Figure S1.** Protocol for ArLV1-detection in plant material.

Schematic representation of the protocol to detect ArLV1 in plant samples using RT-qPCR.

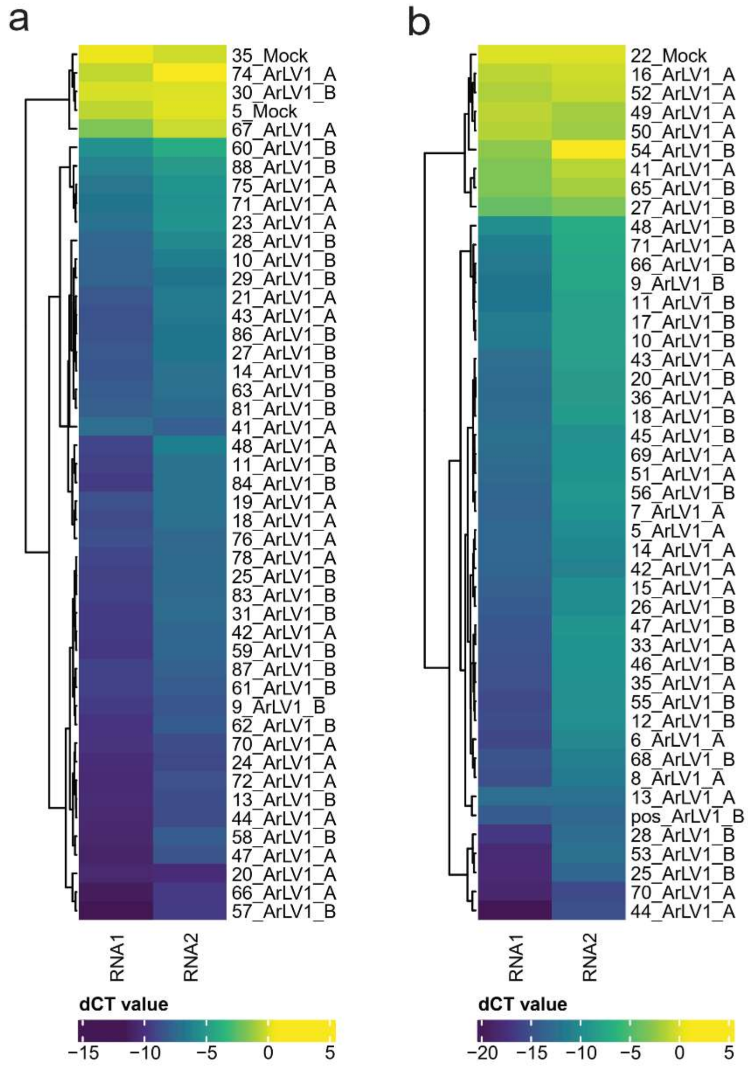


**Figure S2.** *Arabidopsis* inoculation efficiency.

Samples from newly-formed leaves after inoculation of healthy Col-0 plants with ArLV1_A (Wageningen isolate), ArLV1_B (Utrecht isolate) or mock inoculum (Mock). Sample numbers represent the individually sampled plants from two separate experiments (**a** and **b**) for which plant material was tested by qPCR for RNA1 and RNA2 of ArLV1 and dCT was calculated by subtraction from the average mock sample. dCT-values of RNA1 and RNA2 cluster into two main groups (positive and negative**;** *k-means* clustering). In total, 79 of the 90 inoculated plants tested positive for ArLV1, which represents an inoculation efficiency of 88%.

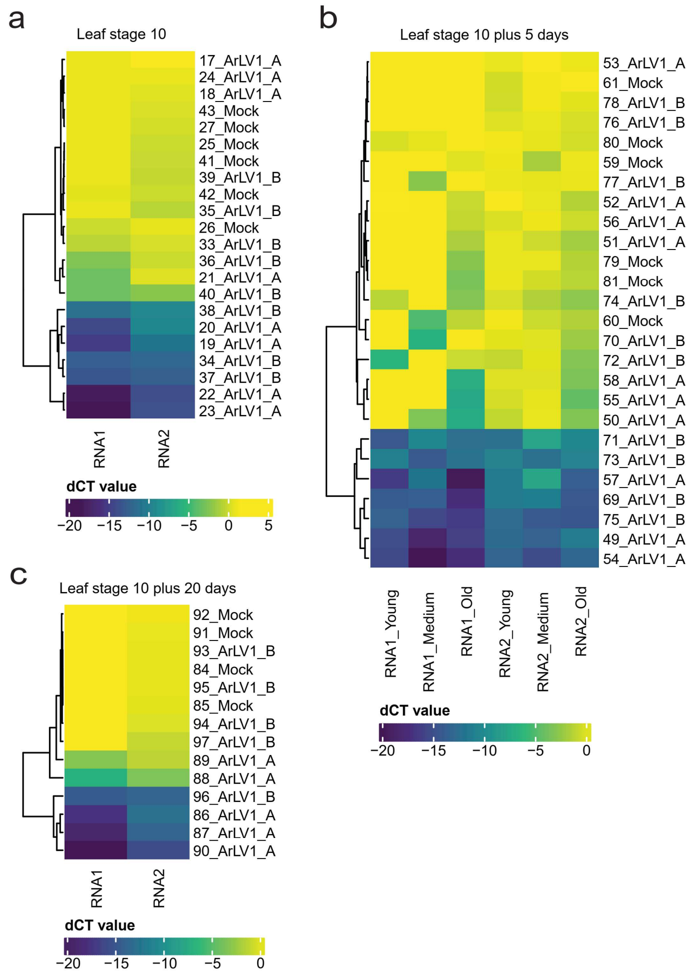


**Figure S3.** *Arabidopsis* seed transmission of ArLV1.

Leaf samples were harvested from individual plants at leaf stage 10 (**a**), leaf stage 10 plus an additional 5 days (**b**), or at leaf stage 10 plus an additional 20 days (**c**). Sample numbers represent the individually sampled plants. RNA extraction and cDNA synthesis were performed according to the described methods. Seeds were acquired from parent Col-0 plants positive for ArLV1-infection (ArLV1_A or ArLV1_B) or healthy naïve parent plants (Mock). Plant material was tested by qRT-PCR for RNA1 and RNA2 of ArLV1 and dCT was calculated by subtraction from the average of the mock samples, so that negative samples are around a dCT of 0. Delta CT-values of RNA1 and RNA2 were divided into two main groups (positive and negative) by *k-means* clustering. In total, 18 of the 46 plants grown from seed produced by infected parent plants tested positive for ArLV1, which represents a seed transmission of 39%.

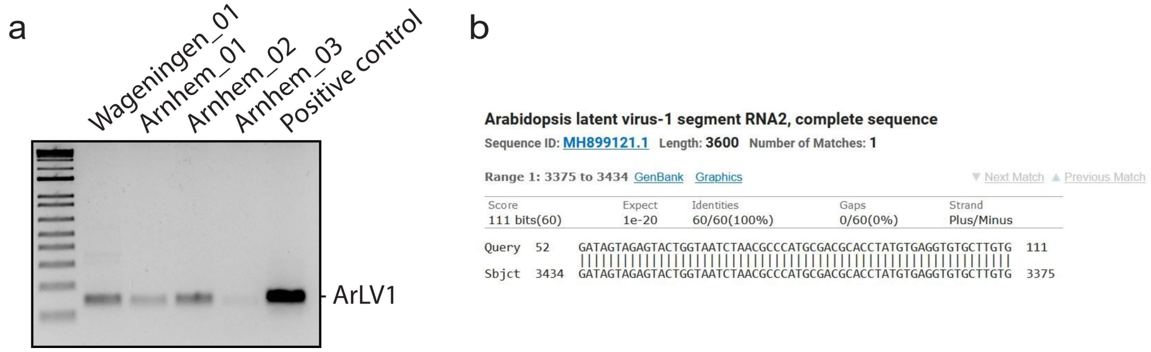


**Figure S4.** ArLV1 RNA2 PCR fragments of wild collected *Arabidopsis*. (**a**) Gel electrophoreses image of PCR fragments of RNA2 derived from plants sampled on two locations in The Netherlands: Wageningen (51°58'53.2"N 5°40'10.5"E) and Arnhem (N51° 59'09.0"N 5°53'52.0"E). (**b**) Sequenced PCR fragment of 'Arnhem_02' from (a).

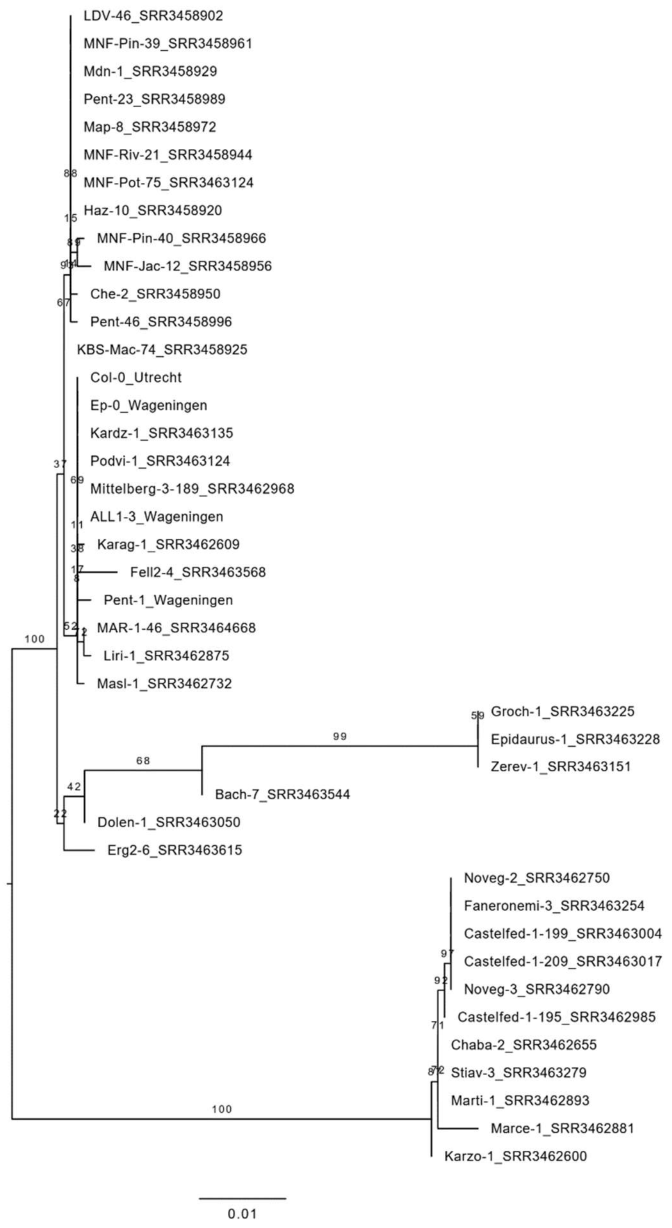


**Figure S5.** Phylogenetic tree of the Pro-Pol region from ArLV1 RNA1. Branch labels indicate bootstrap values. Note, this is an unrooted tree. Nucleotide sequences are from 38 NCBI SRA datasets, with a full coverage of RNA1 and RNA2 supplemented with four sequences from different *Arabidopsis* ecotypes from Wageningen and Utrecht (ALL1-3_Wageningen (=ArLV1_A), Pent-1_Wageningen, Ep-0_Wageningen and Col-0_Utrecht (=ArLV1_B)). Branch lengths (scale) represent nucleotide substitutions per site.

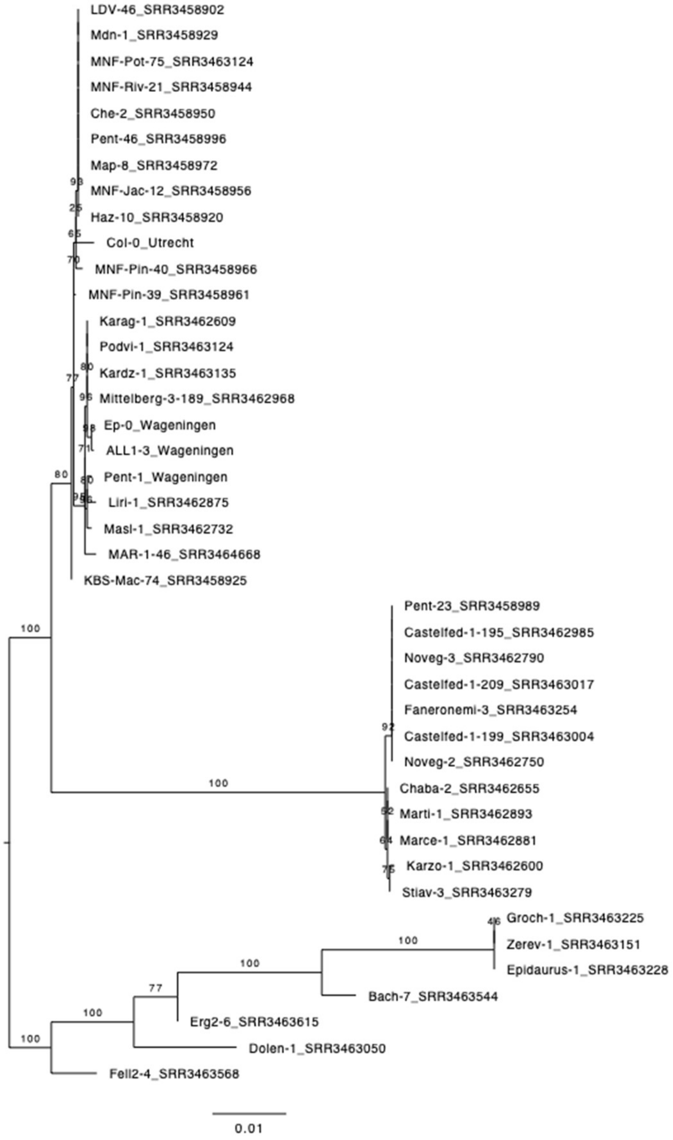


**Figure S6.** Phylogenetic tree of the full ORF from ArLV1 RNA2. Branch labels indicate bootstrap values. Note, this is an unrooted tree. Nucleotide sequences are from 38 NCBI SRA datasets, with a full coverage of RNA1 and RNA2 supplemented with four sequences from different *Arabidopsis* ecotypes from Wageningen and Utrecht (ALL1-3_Wageningen (=ArLV1_A), Pent-1_Wageningen, Ep-0_Wageningen and Col-0_Utrecht (=ArLV1_B)). Branch lengths (scale) represent nucleotide substitutions per site.

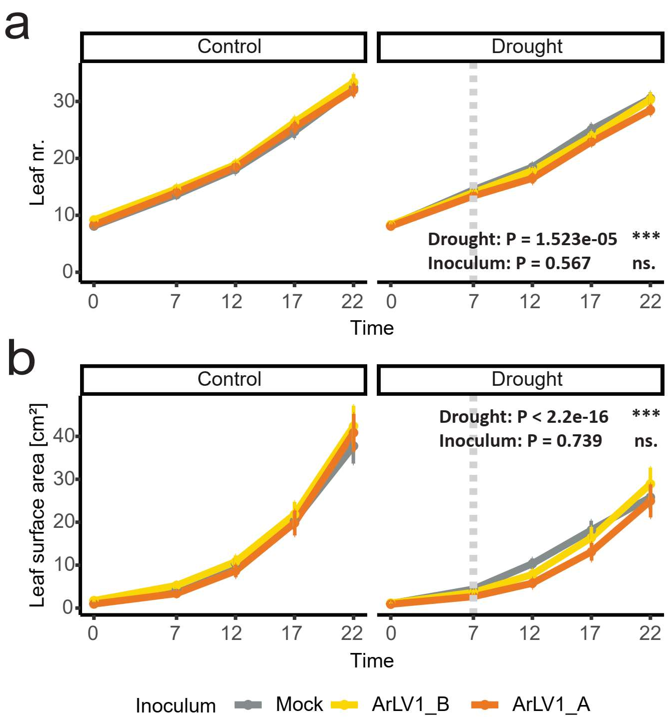


**Figure S7.** Leaf number and leaf surface area of plants subjected to ArLV1-inoculation. Phenotypic observations after inoculation of healthy Col-0 plants with ArLV1_A (Wageningen isolate), ArLV1_B (Wageningen isolate) or mock inoculum (Mock). Leaf number (a) and leaf surface area (b) were measured at different timepoints (days) after inoculation with ArLV1 (started Time = 0 days) and after drought was applied (started at Time = 7 days, indicated by the vertical dotted bar). The experiment was repeated twice with similar results. Both repeats were included in the statistical analysis using mixed linear models with repeats as a random variable and an alpha of 0.05.

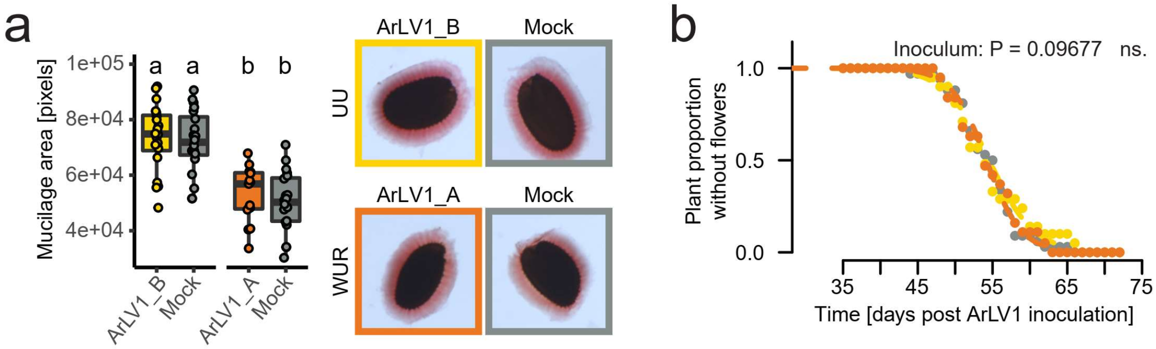


**Figure S8.** Seed mucilage layer and flowering time are not affected by ArLV1. (**a**) Quantification of seed pectin mucilage area after staining with ruthenium red (left) with representative pictures (right). Seeds of plants infected with ArLV1_A (Wageningen isolate: WUR) or ArLV1_B (Utrecht isolate: UU), and their respective mock seed batches, were derived from plants grown in the different labs. Different letters (a, b) represent significant differences (P< 0.05). Boxplots represent the minimum, first quantile, median, third quantile and maximum and all individual measurements (circles; n>17). (**b**) Fractions of plants that do not flower. The experiment was repeated twice with similar results. Both repeats were included in the statistical analysis using generalized mixed linear models with repeats as a random variable and an alpha of 0.05. The p-values of Inoculum represents the effect of inoculation with either ArLV1_A (orange), ArLV1_B (yellow) or Mock (grey).

**Table S1.** *Arabidopsis* SRA-datasets included in the geographical analysis. SRA datasets were downloaded from the NCBI Sequence Read Archive (https://www.ncbi.nlm.nih.gov/sra). The ArLV1-clade from the phylogenetic analysis is given as well as the country of origin of the Arabidopsis line (BUL: Bulgaria, FRA: France, GER: Germany, GRC: Greece, ITA: Italy, NL: The Netherlands, RUS: Russia, SPA: Spain, SVK: Slovakia, USA: United States of America).

| | Arabidopsis genotype | Origin | ArLV1 clade | Origin country |
|---|---|---|---|---|
| 1 | Bach-7 | SRR3463544 | 3 | GER |
| 2 | Castelfed-1-195 | SRR3462985 | 2 | ITA |
| 3 | Castelfed-1-199 | SRR3463004 | 2 | ITA |
| 4 | Castelfed-3-209 | SRR3463017 | 2 | ITA |
| 5 | Chaba-2 | SRR3462655 | 2 | RUS |
| 6 | Dolen-1 | SRR3463050 | 3 | BUL |
| 7 | Epidauros-1 | SRR3463228 | 3 | GRC |
| 8 | Erg2-6 | SRR3463615 | 3 | GER |
| 9 | Faneronemi-3 | SRR3463254 | 2 | GRC |
| 10 | Groch-1 | SRR3463225 | 3 | BUL |
| 11 | Haz-10 | SRR3458920 | 1 | USA |
| 12 | Karag-1 | SRR3462609 | 1 | RUS |
| 13 | Kardz-1 | SRR3463135 | 1 | BUL |
| 14 | KBS-Mac-74 | SRR3458925 | 1 | USA |
| 15 | Krazo-1 | SRR3462600 | 2 | RUS |
| 16 | LDV-46 | SRR3458902 | 1 | FRA |
| 18 | Map-8 | SRR3458972 | 1 | USA |
| 20 | Marce-1 | SRR3462881 | 2 | ITA |
| 21 | Marti-1 | SRR3462893 | 2 | ITA |
| 22 | Masl-1 | SRR3462732 | 1 | RUS |
| 23 | Mdn-1 | SRR3458929 | 1 | USA |
| 24 | Mitterberg-3-189 | SRR3462968 | 1 | ITA |
| 25 | MNF-Che-2 | SRR3458950 | 1 | USA |
| 26 | MNF-Jac-12 | SRR3458956 | 1 | USA |
| 27 | MNF-Pin-39 | SRR3458961 | 1 | USA |
| 28 | MNF-Pin-40 | SRR3458966 | 1 | USA |
| 29 | MNF-Pot-75 | SRR3458937 | 1 | USA |
| 30 | MNF-Riv-21 | SRR3458944 | 1 | USA |
| 31 | Noveg-2 | SRR3462750 | 2 | RUS |
| 32 | Noveg-3 | SRR3462790 | 2 | RUS |
| 33 | Pent-23 | SRR3458989 | 1 | USA |
| 34 | Pent-46 | SRR3458996 | 1 | USA |
| 35 | Podvi-1 | SRR3463124 | 1 | BUL |
| 36 | Stiav-3 | SRR3463279 | 2 | SVK |
| 37 | Wild plant | Wageningen region | Not sequenced | NL |
| 38 | Wild plant | Arnhem region | Not sequenced | NL |
| 39 | Wild plant | Madrid region | Not sequenced | SPA |
| 40 | ALL1-3 | Wageningen lab | 1 | NL |
| 41 | Pent-1 | Wageningen lab | 1 | NL |
| 42 | Ep-0 | Wageningen lab | 1 | NL |
| 43 | Col-0 | Utrecht lab | 1 | NL |

**Methods S1** RNA-Seq approaches and plant growth, treatment and sampling conditions

Dataset Wageningen University and Research

Study design, sampling, data cleaning and read mapping were performed as described earlier (Kloth *et al.*, 2016).

Dataset Utrecht

Growth conditions and treatments are largely as described in Morales *et al.,* (2022). *Arabidopsis* Col-0 seeds were sown on a moist soil:perlite mix 1:2 (Primasta BV, Asten, The Netherlands) and stratified in dark conditions for five days at 4 $^{o}$C, after which germination was induced at either 21 $^{o}$C or 27 $^{o}$C (for high temperature treated plants) at a 8 h light, 16 h dark short-day regime. Seedlings were transplanted to individual Jiffy 7c coco-pellets (Jiffy Products International BV, Zwijndrecht, The Netherlands), when they reached the 2-leaf stage (about 1.5 week after sowing) and further cultivated under identical short-day conditions. Hoagland solution was applied at two, five and seven days after transplanting (10 mL, 20 mL and 10 mL per plant respectively). Plants were watered every two days unless subjected to drought or submergence treatment. Plants were randomized every 2-3 days up to 10-leaf stage. At 10-leaf stage, plants were subjected to different stress conditions: mild drought, high temperature, high temperature combined with mild drought, 5-day submergence in light, post submergence followed by recovery, and post submergence combined with mild drought.

Leaf material was harvested at 0, 5 and 10 days after the treatments started and all samples were collected at 2 hours after the photoperiod began. For each sample, five or six plants were selected and two young leaves (leaves 7, 8, 9 and 10 are defined as young leaves, counting from the oldest leaf at 10-leaf stage) from each plant were detached and pooled together as a biological replicate. Leaf pools were immediately snap-frozen with liquid nitrogen. Frozen leaves were ground to a fine powder and processed to total RNA extraction using the RNeasy kit (Qiagen, Germany) following the manufacturers protocol. Samples were finally diluted to 25 ug/ul with a total volume of 60 uL using DEPC-treated water.

Quality control, library construction and Illumina sequencing were carried out by MACROGEN (Amsterdam, The Netherlands). Total RNA integrity and pureness were checked using an Agilent Technologies 2100 Bioanalyzer (Agilent, Santa Clara). Library preparations were performed based on the TruSeq stranded mRNA protocol (Illumina, San Diego). The constructed libraries were then sequenced by Illumina Novaseq6000 sequencer with 150 bp pair-end reading and finally provided with FASTQ files output.

Data cleaning, including adaptor trimming and reads filtering was conducted using *CUTADAPT* (Martin, 2011) under Linux operation system. The Truseq adaptor sequences "AGATCGGAAGAGCACACGTCTGAACTCCAGTCA" and "AGATCGGAAGAGCGTCGTGTAGGGAAAGAGTGT" were trimmed with a maximum error rate of 0.07. Reads filtering was based on the criterion of the minimum cutoff length of 30 bp and quality score of 20. This brought at least 61 million clean pair-end reads per sample. The trimmed and filtered reads were then assembled and mapped onto *Arabidopsis* transcriptome (Araport10) using *KALLISTO* (Bray *et al.*, 2016).

**Methods S2** Transcriptomic analysis of Arabidopsis Col-0 accession

For details of plant growth conditions and treatments used for the Utrecht transcriptome dataset, see Supplemental Information Methods S1. In short, 58 samples of young leaves from Arabidopsis Col-0 at different timepoints and under different abiotic stress treatments (Morales *et al.*, 2022) were harvested. Total RNA was extracted, a library was made and sequenced using the Truseq stranded mRNA protocol and an Illumina Novaseq6000. Before analysis, the transcripts per million (TPM) values were filtered and transformed in R version 4.0.3 x64. Next, Arabidopsis gene expression was filtered for read detection of $\log_2 > 2$ in all samples. This filter resulted in 6087 detected genes (out of 27655 protein coding genes in the assembly). Subsequently, the TPM values were transformed by

$$TPM_{log,i,j} = \log_2(TPM_{i,j} + 1)$$

where $TPM_{log}$ was the $\log_2$-normalized TPM value of gene *i* (one out of 6048) and sample *j* (one out of 58 samples).

For correlation analysis, a ratio was also calculated with the mean of the TPM, by

$$TPM_{ratio,i,j} = \log_2(\frac{TPM_{i,j}}{\overline{TPM_i}})$$

where $TPM_{ratio}$ was the $\log_2$ of the TPM value of gene *i* (one out of 6087) and sample *j* (one out of 58 samples), divided by the average TPM value over all samples for gene *i*. To understand the sources of variance in the expression data, principal component analyses were done with the *prcomp* function in R with the parameter *scale. = TRUE* on the $TPM_{ratio}$-transformed expression data. Likewise, correlation matrices were calculated on the $TPM_{ratio}$-transformed expression data with *cor* and the *heatmap* function in R. We selected seven samples taken at timepoint 5 under no applied abiotic stress, as three samples of this selection contained relatively few reads mapping to ArLV1 (0.01-9.56%) and four samples contained high amounts of ArLV1 (78.94-90.08%).

Statistical analysis was done in R version 4.1.2 x64 using the $\log_2$-normalized TPM values in linear models ran for timepoint 5 for the control samples containing a high ArLV1 content when compared to the control samples with a low ArLV1 content at the same timepoint. The obtained significances were corrected using a Benjamini-Hochberg adjustment for multiple testing (provided by the *prcomp* function). Visualization of all genes in a volcanoplot was done with the ggplot2 package (Wickham, 2016).

**Methods S3** Phenotypic parameters

At different timepoints (-7; the day of inoculation), 0 (the day were the drought treatment started, if applicable), 5, 10, and 15 days), several phenotypic characteristics of the plants were measured. Leaf number was counted visually, and weight was measured including the pot and coco-pellet (Jiffy, Zwijndrecht) to estimate soil drying. Leaf chlorophyll content was measured using a CCM-300 chlorophyll content meter (Opti-Sciences, USA) on the 8$^{th}$ true leaf. Leaf surface area was measured by taking a picture of all plants from above (perpendicular to the rosette) and creating a color threshold in ImageJ (Schneider *et al.*, 2012). Drought was applied by withholding the application of water. See also Morales *et al.,* (2022). Wilting timepoints were scored when plants lost turgor, and flowering was scored when the first flower was open. Statistical analyses and visualizations were performed using R version 4.1.2. For normally distributed data, one way ANOVA's were used with a Tukey's posthoc test. For nested data with a logistic regression (flowering point and wilting point), generalized linear mixed effects regression models (*glmer*) were implemented. For nested data with a normal regression (leaf number, weight, chlorophyll content and leaf surface area), linear mixed effects regression models (*lmer*) were implemented. The R packages lme4 (Bates *et al.*, 2015) and emmeans (Lenth, 2022) were used for these analyses.

### Seed mucilage assays

Seeds obtained from ArLV1-positive and negative plants from Wageningen (ArLV1_A) and Utrecht (ArLV1_B) were immersed in MilliQ for three minutes. Aqueous ruthenium red solution was added to a final dilution of 0.2% (w/v) and seeds were incubated for 15 minutes. Seeds were then washed three times in MilliQ before pictures were taken on a Zeiss Axiozoom v16. Pictures were analyzed using ImageJ to determine the size of the seed and the mucilage layer surrounding it.
